# Structural insights into hormone recognition by the human glucose-dependent insulinotropic polypeptide receptor

**DOI:** 10.1101/2021.03.18.436101

**Authors:** Fenghui Zhao, Chao Zhang, Qingtong Zhou, Kaini Hang, Xinyu Zou, Yan Chen, Fan Wu, Qidi Rao, Antao Dai, Wanchao Yin, Dan-Dan Shen, Yan Zhang, Tian Xia, Raymond C. Stevens, H. Eric Xu, Dehua Yang, Lihua Zhao, Ming-Wei Wang

## Abstract

Glucose-dependent insulinotropic polypeptide (GIP) is a peptide hormone that exerts crucial metabolic functions by binding and activating its cognate receptor, GIPR. As an important therapeutic target, GIPR has been subjected to intensive structural studies without success. Here, we report the cryo-EM structure of the human GIPR in complex with GIP and a G_s_ heterotrimer at a global resolution of 2.9 Å. GIP adopts a single straight helix with its N terminus dipped into the receptor transmembrane domain (TMD), while the C-terminus is closely associated with the extracellular domain and extracellular loop 1. GIPR employs conserved residues in the lower half of the TMD pocket to recognize the common segments shared by GIP homologous peptides, while uses non-conserved residues in the upper half of the TMD pocket to interact with residues specific for GIP. These results provide a structural framework of hormone recognition and GIPR activation.

## Introduction

Glucose-dependent insulinotropic polypeptide (GIP) is a 42-amino acid peptide hormone that plays crucial role in glucose regulation and fatty acid metabolism. In response to food intake, GIP is secreted by intestinal K cells to enhance insulin secretion and peripheral fatty acid uptake (*1*), as well as a number of neuronal effects (*2*). The pleiotropic functions of GIP is mediated by its cognate receptor (GIPR), a member of class B1 G protein-coupled receptors (GPCRs) that also include glucagon receptor (GCGR) and glucagon-like peptide-1 (GLP-1R). GIPR, together with GCGR and GLP-1R, form the central endocrine network in regulating insulin sensitivity and energy homeostasis, and they are validated drug targets (*3–5*). Intensive efforts were made in drug discovery targeting these receptors (*6*). A number of GLP-1R selective ligands have been developed successfully to treat type 2 diabetes and obesity. Encouragingly, peptide ligands that bind both GIPR and GLP-1R show better clinical efficacy than the GLP-1R agonist alone. As such, GIPR has emerged as a hot target pursued by pharmaceutical research community.

GIPR contains a large extracellular domain (ECD) and a 7-transmembrane domain (TMD). Both are involved in ligand recognition and receptor activation (*7–9*). Cryo-electron microscopy (cryo-EM) structures of GCGR and GLP-1R, as well as several other class B1 GPCRs have been solved, providing a general mechanism of two-domain model for peptide recognition and receptor activation. However, GIP displays an exquisite sequence specificity towards GIPR as it does not bind to other class B1 GPCRs. However, the efforts to understand the ligand selectivity by GIPR have been hampered by technical difficulties in expression and stabilization of the liganded GIPR complexes for structural studies. We have overcome such challenges and determined a high-resolution (2.9 Å) structure of the human GIPR in complex with the stimulatory G protein (G_s_) using single-particle cryo-EM approach in conjunction with NanoBiT strategy (*10*). Together with functional studies, our results demonstrate several unique structural features that distinguish GIPR from other members of the glucagon subfamily of class B1 GPCRs and provide an important template for rational design of GIPR agonists for therapeutic development.

## Results

### Structure determination

To prepare a high quality human GIPR–G_s_ complex, we overcame several technical obstacles to enhance the expression level and protein stability by adding a double tag of maltose binding protein at the C terminus and a BRIL fusion protein at the N terminus (Fig. S1A), as well as employing the NanoBiT tethering strategy (*10–12*) (Fig. S1A, B). To solve the GIP_1-42_–GIPR–G_s_ structure, we further introduced one mutation (T345F) to stabilize the assembly of complex (Fig. S1C, D). This mutation does not affect the ligand binding or potency of GIP_1-42_ in cAMP accumulation assay (Fig. S1G, H). Large-scale purification was followed and the GIP_1-42_–GIPR–G_s_ complexes were collected by size-exclusion chromatography (SEC) for cryo-EM studies (Fig. S1E, F). The activity of the modified GIPR construct was confirmed by cAMP accumulation assay showing a similar response as that of the wild-type (WT; Fig. S1G).

The GIP_1-42_–GIPR–G_s_ complexes were imaged using a Titan Krios equipped with a Gatan K3 Summit direct electron detector (Fig. S2). 2D classification showed a clear secondary structure feature and random distribution of the particles. Different directions of the particles enabled a high-resolution cryo-EM map reconstruction (Fig. S2B). A total of 295,021 particles were selected after 3D refinement and polishing, leading to an overall resolution of 2.9 Å (Fig. S2C, D).

### Overall structure

Apart from the α-helical domain (AHD) of Gα_s_ which is flexible in most cryo-EM GPCR–G protein complex structures, the bound GIP_1-42_, GIPR and G_s_ were well defined in the EM density maps (Fig. 1, Fig. S3). Except for the ECD, side-chains of the majority of amino acid residues are well resolved in all protein components. The final model contains 30 GIP_1-42_ residues, the Gαβγ subunits of G_s_, and the GIPR residues from Q30^ECD^ to S415^8.66b^ (class B GPCR numbering in superscript) (*13*), with six amino acid residues missing at helix 8. Owing to the high resolution map, notable conformation difference from GCGR (*14*) or GLP-1R (*15*) was observed in the ECL1.

**Fig. 1.**
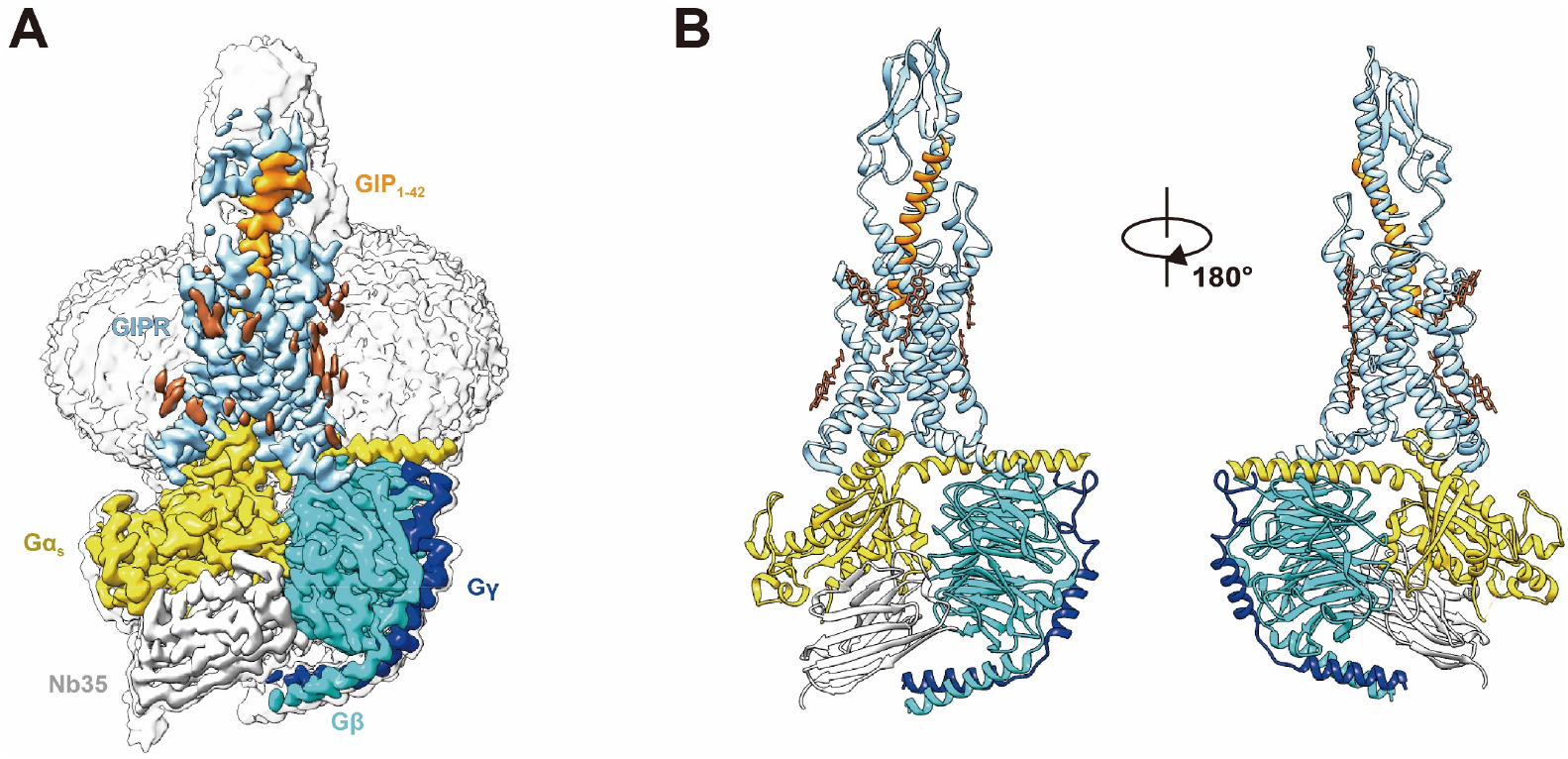
Cryo-EM structure of the GIP_1–42_–GIPR–G_s_ complex. (**A**), Cut-through view of the cryo-EM density map that illustrates the GIP_1–42_–GIPR–G_s_ complex and the disc-shaped micelle. The unsharpened cryo-EM density map at the 0.07 threshold shown as light gray surface indicates a micelle diameter of 11 nm. The colored cryo-EM density map is shown at the 0.16 threshold. (**B**), Model of the complex as a cartoon, with GIP_1–42_ as helix in orange. The receptor is shown in light sky blue, Gα_s_ in yellow, Gβ subunit in cyan, Gγ subunit in navy blue and Nb35 in gray.

Similar to other class B1 GPCR–G_s_ complexes, the TM6 of GIPR shows a sharp kink in the middle and TM7 displays an outward movement. Like PTH1R–G_s_ and CRF1R–G_s_ cryo-EM structures (*16, 17*), the TMD of GIPR is surrounded by annular detergent micelle, with a diameter of 12 nm thereby mimicking the lipid bilayer morphology (Fig. 1). In addition, we also observed several cholesterols molecules in the cryo-EM map.

### Ligand recognition

In the complex, GIP adopts a single continuous helix that penetrates into the TMD core through its N-terminal half (residues 1 to 15), while the C-terminal half (residues 16 to 30) is recognized by the ECD and ECL1 (Fig. 2A-C). Y1^P^ (P indicates that the residue belongs to the peptide ligand) of GIP points to TMs 2-3, forms hydrogen bonds with R190^2.67b^ and Q224^3.37b^, and makes hydrophobic contacts with V227^3.40b^ and W296^5.36b^. This observation received support of the mutagenesis study, where mutant W296A decreased GIP potency by 50-fold (Fig. 2D), and mutants R190A and Q224A diminished the potency of GIP by 71- and 5-fold, respectively, as reported in a previous report (*18*). N-terminal truncation of either Y1^P^ or both Y1^P^ and A2^P^ led to reduced low efficacy or loss of activity (*19, 20*), highlighting a crucial role of Y1^P^. E3^P^, D9^P^ and D15^P^ are three negatively charged residues in the N-terminal half of GIP and form salt bridges with R183^2.60b^, R370^7.35b^ and R289^ECL2^, respectively. Removal of these salt bridges by alanine substitution at either R183^2.60b^ (*18*) or R370^7.35b^ (Fig. 2D) greatly reduced GIP potency (by 76- and 55-fold, respectively), whereas the effect on mutant R289A was mild (6-fold, Fig. 2D). Polar interactions also occurred between S8^P^ and N290^ECL2^ as well as Y10^P^ and Q138^1.40b^. The GIP–TMD interface was further stabilized by a complementary nonpolar network involving TM1 (L134^1.36b^, L137^1.39b^ and Y141^1.43b^) and TM7 (L374^7.39b^ and I378^7.43b^) via A2^P^, F6^P^ and Y10^P^ of GIP (Fig. 2C), in line with decreased ligand potencies observed in Y141A (by 103-fold), L374A (by 41-fold) and I378A (by 8-fold) mutants (Fig. 2D).

**Fig. 2.**
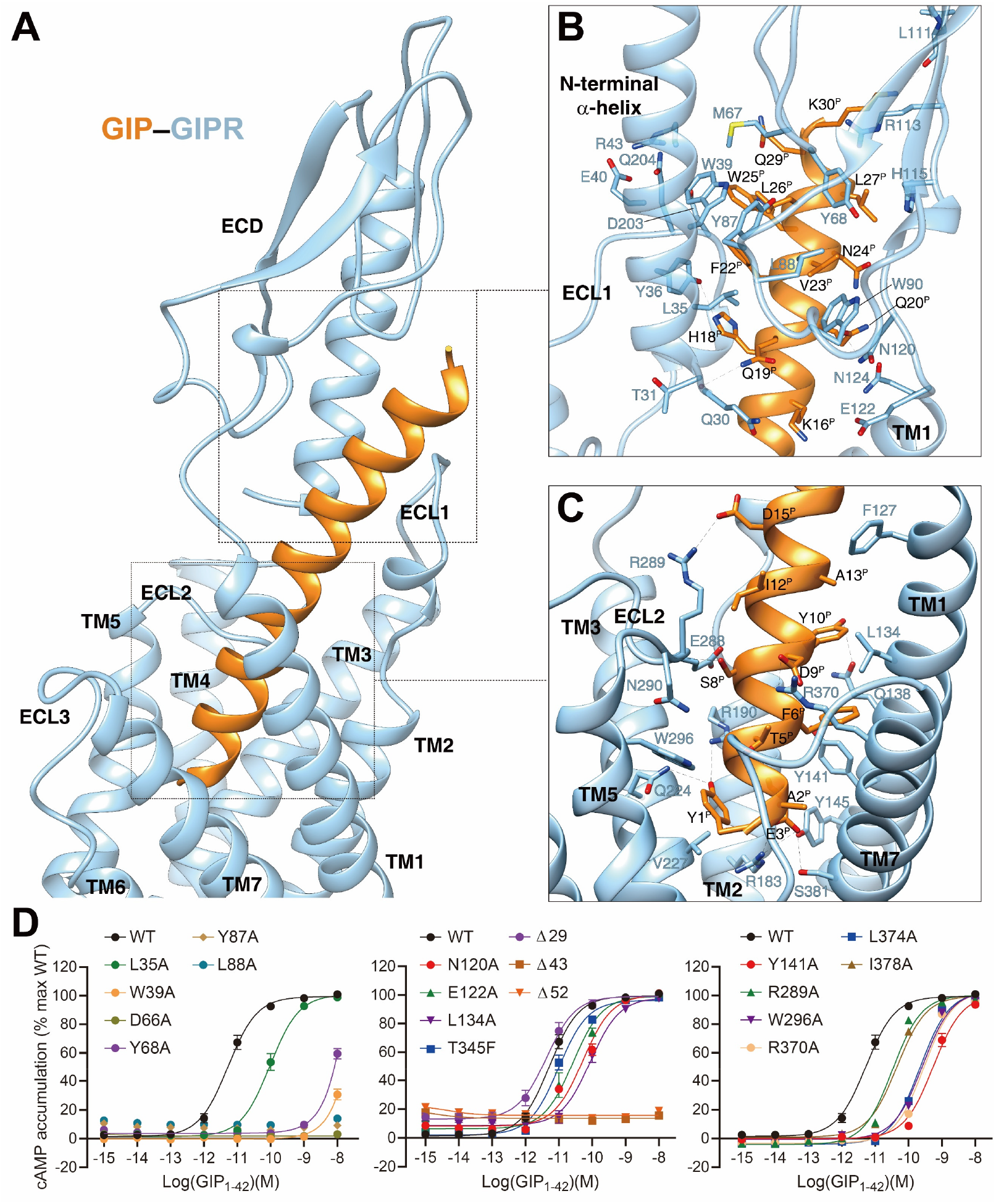
Molecular recognition of GIP by GIPR. (**A**), the binding mode of GIP (orange) with GIPR (light sky blue), showing that the N-terminal half of GIP penetrates into a pocket formed by all TM helices except TM4, ECL2 and ECL3, whereas the C-terminal half is recognized by ECD, ECL1 and TM1. (**B-C**), close-up views of the interactions between GIP and GIPR. (**D**), Signaling profiles of GIPR mutants. cAMP accumulation in wild-type (WT) and single-point mutated GIPR expressing in HEK 293T cells. Signals were normalized to the maximum response of the WT and dose-response curves were analyzed using a three-parameter logistic equation. All data were generated and graphed as means ± S.E.M. of at least three independent experiments, conducted in quadruplicate. ∆, truncated residues.

The C-terminal half of GIP was clasped by the GIPR ECD, closely resembling the crystal structure of GIP–GIPR ECD (PDB code: 2QKH) (*21*). Specifically, three adjacent amino acids (H18^P^, Q19^P^ and Q20^P^) form multiple hydrogen bonds with the side-chains of Q30 and Y36 at the N-terminal α-helix of ECD, as well as N120 and E122 at the hinge region between ECD and TM1. The hydrophobic residues of GIP (F22^P^, V23^P^, L26^P^ and L27^P^) occupy a complementary binding groove of the GIPR ECD, consisting of a series of hydrophobic residues (L35, Y36, W39, M67, Y68, Y87, L88, P89 and W90). Alanine substitutions in W39, D66 and Y68 significantly reduced the potency of GIP (Fig. 2D). Notably, the cryo-EM map suggests that the ECL1 stands upwards to approach the N-terminal α-helix of ECD and forms hydrogen bonds with the side-chains of R43 and Y36 (Fig. 2A, B), resulting in a close contact between TMD and ECD for GIP-bound GIPR (interface area = 571 Å^2^), significantly larger than that of GLP-1 bound GLP-1R (362 Å^2^), reinforcing the importance of ECD in GIP recognition.

### Receptor activation

GIPR shares ~50% sequence similarity with GCGR, especially in the TMD region (75%), thus GCGR structures published previously provide a good template for the present study (Fig. S4) (*14, 22–25*). It was found the TMD of activated GIPR exhibits a conformation similar to that of GCGR activated by glucagon or ZP3780 (Cα RMSD = 1.2 and 0.7 Å, respectively) (*14, 22*) and distinct from that of GCGR bound by the negative allosteric modulator NNC0640 or partial agonist NNC1702 (Cα RMSD = 4.0 and 3.9 Å, respectively) (*26*). Facilitated by Gly^7.50b^ located in the middle of TM7, the extracellular half of TM7 bends towards TM6 by 8.0 Å (measured by Cα atom of Gly^7.32b^)(Fig. S4). This feature and the outward movement of ECL3 expanded the ligand binding pocket. Meanwhile, the extracellular tip of TM1 was extended by one turn and moved inward by 8.0 Å (measured by Cα atom of the residues at 1.30b)(Fig. S4). Together with the raised ECL1, these conformational changes stabilized ligand binding.

In the intracellular side, the sharp kink in the middle of TM6 led to an outward movement of its intracellular portion measured by Cα atom of R336^6.35b^ (18.9 Å, similar to that of other G_s_-coupled class B1 receptors). This was accompanied by the movement of the intracellular tip of TM5 towards TM6 by 7.6 Å (measured by Cα atom of the residues at 5.67b), thereby creating an intracellular cavity for G protein coupling (Fig. S4).

### G protein coupling

In our model, G_s_ protein is anchored by the α5 helix of Gα_s_ (GαH5), thereby fitting to the cytoplasmic cavity formed by TMs 3, 5 and 6, intracellular loops (ICLs) 1-2 and H8 (Fig. 3). In general, the GIPR–G_s_ complex shows a similar receptor–G protein interface as other reported class B1 receptor structures such as GLP-1R (*27*), GLP-2R (*12*), GCGR (*14*), PTH1R (*16*), SCTR (secretin receptor) (*28*) and GHRHR (*11*), suggesting a common G protein signaling mechanism (Fig. 3A). The hydrophobic residues at the C-terminal of GαH5 (L388^GαH5^, Y391^GαH5^, L393^GαH5^ and L394^GαH5^) insert into a small hydrophobic pocket formed by Y240^3.53b^, L241^3.54b^, L244^3.57b^, L245^3.58b^, I317^5.58b^, I320^5.60b^, L321^5.61b^ and L325^5.65b^ (Fig. 3B). The side-chain of R338^6.37b^ points to Gα_s_ and makes one hydrogen bonds with L394^GαH5^. Of note is the interaction between R380^GαH5^ and ICL2 resulting in five hydrogen bonds with the backbone atoms of L245^3.58b^, V246^3.59b^, L247^3.60b^ and V248^ICL2^, significantly more than that observed in GLP-1R, SCTR or GCGR (Fig. 3C). The polar residues in ICL2 (S251^ICL2^ and E253^ICL2^) produce two hydrogen bonds with K34 and Q35 of Gα_s_, while H8 forms several hydrogen bonds with ICL1, then contacts with Gβ (E398^8.49b^-R164^ICL1^-D312^Gβ^, E402^8.53b^-R164^ICL1^-D312^Gβ^) (Fig. 3D). Together, these specific interactions contribute to the Gs coupling specificity of GIPR.

**Fig. 3.**
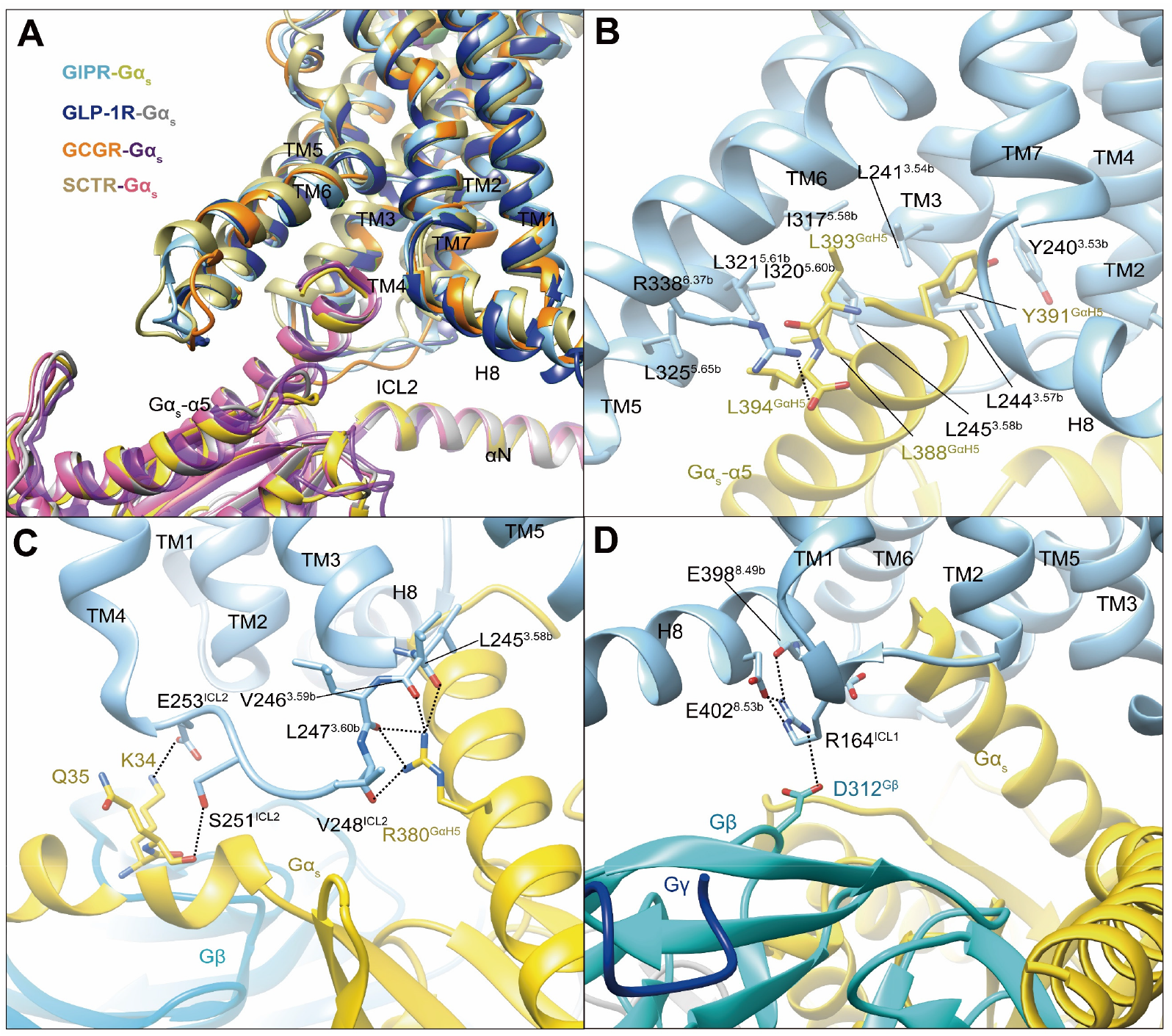
G protein coupling of GIPR. (**A**), Comparison of G protein coupling among GIPR, GLP-1R (*27*), GCGR (*14*) and SCTR (*28*). The Gα_s_ α5-helix of the Gα_s_ Ras-like domain inserts into an intracellular crevice of GIPR TMD. (**B**), Interaction between GIPR and the C-terminus of Gα_s_. (**C**), Polar interactions between ICL2 and Gα_s_. (**D**), Polar interactions between H8 and ICL1 of the GIPR and Gβ. The GIP_1–42_–GIPR–Gα_s_ structure is colored light sky blue (GIPR), gold (Gα_s_) and cyan (Gβ). Residues involved in interactions are shown as sticks. Polar interactions are shown as black dashed lines.

### Ligand specificity

GIP, GLP-1 and glucagon are three important metabolic hormones exerting distinct functions in glucose homeostasis, in spite of high degrees of sequence similarity. Superimposing the TMD of GIP-bound GIPR with that of GLP-1-bound GLP-1R (*27*) or glucagon-bound GCGR (*14*) displays a similar ligand-binding pocket and the three peptides all adopt a single continuous helix, with the N-terminus penetrating to the TMD core to the same depth, while the C-terminus anchors the ECD and ECL1 in a receptor-specific manner (Fig. 4). Notably, the ECL1 of GIPR stands upwards in line with TMs 2 and 3, and moves towards the TMD core by 5~7 Å. Such a movement, together with a α-helical extension in TM1 by six residues, allows GIP to shift to TM1 by 2.7 and 3.3 Å (measured by Cα atom of L27^P^) relative to GLP-1 (*27*) and glucagon (*14*), respectively (Fig. S5).

**Fig. 4.**
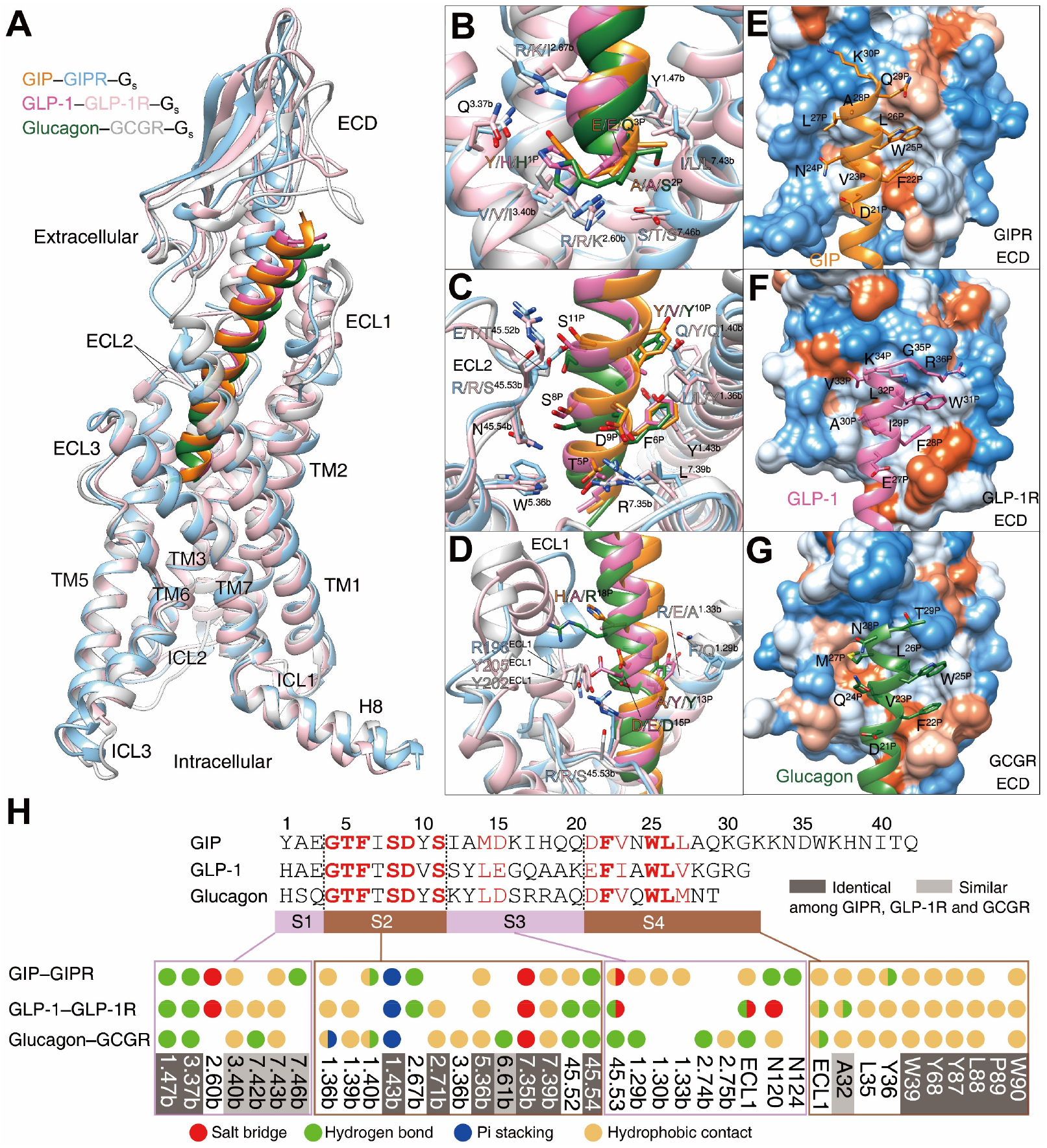
Ligand specificity among GIPR, GLP-1R and GCGR. (**A**), Comparison of the overall structures of GIP_1–42_–GIPR–G_s_, GLP-1–GLP-1R–G_s_ (*27*) and glucagon–GCGR–G_s_ complexes (*14*). G proteins are omitted for clarity. (**B-D**), Close-up views of the interaction between TMD and peptide. Based on sequence similarity, the peptides are divided into four segments: N-terminus (residues 1–3, **B**), segment 2 (residues 4–11, **C**), segment 3 (residues 12–20, **D**) and the C-terminus (residues 21 to the end, **E-G**), where segments 2 and 4 are highly conserved among GIP, GLP-1 and glucagon. Residues are numbered based on GIP for peptides, and labeled with class B GPCR numbering in superscript for receptors (*13*). (**E-G**), Close-up views of the interface between GIPR ECD and GIP C-terminus (**E**), between GLP-1R and GLP-1 C-terminus (**F**), and between GCGR and glucagon C-terminus (**G**). The ECD is shown in surface representation and colored from dodger blue for the most hydrophilic region, to white, to orange red for the most hydrophobic region. (**H**), Comparison of peptide recognition modes for three receptors, described by fingerprint strings encoding different interaction types of the surrounding residues in each receptor. Peptide residue numbers on the top are shown based on GIP. The ligand-binding pocket residues that are identical or similar across three receptors are highlighted in dark grey and light grey, respectively. Color codes are listed on the bottom.

Based on the sequence similarity, the three peptides can be divided into four segments: two common segments (residues 4-11 and 21 to 30 in GIP) and two unique segments (residues 1-3 and 12-20 in GIP) (Fig. 4H). The N-terminus (residues 1-3) makes massive contacts with the conserved central polar network of class B1 GPCRs including one hydrogen bond with Q^3.37b^ stabilized by the hydrophobic residue at 3.40b; one hydrogen bond with Y^1.47b^ made by the third peptide residue (Fig. 4B, H); residues 4-11 interact with salt bridges of R^7.35b^, pi-stacking of Y^1.43b^, hydrophobic L^2.71b^, W^5.36b^ and L^7.39b^, as well as several hydrogen bonds in ECL2 (Fig. 4C, H); residues 12-20 are divergent and mainly interact with ECLs 1-2 and TMs 1-2 (Fig. 4D, H).

To accommodate varying lengths of side-chains at A^13P^/Y/Y, I^17P^/Q/R, Q^19P^/A/A and Q^20P^/K/Q, both TM1 and ECL1 adjusted their conformations to avoid clashes (Fig. 4D, Fig. S5). For example, ECL1 of GLP-1R is more distant from GLP-1 than that of GIPR from GIP, whereas repulsion of the side-chain of R^18P^ was seen between GCGR and glucagon. Therefore, receptor-specific interaction may reside in this region which precludes the binding of GLP-1 or glucagon to GIPR revealed by MD simulations (Fig. S6). As far as C-terminus is concerned, all three peptides form extensive hydrophobic contacts with the ECD, resulting from the hydrophobic composition of amino acids in both sides (Fig. 4E-H). It appears that GIPR, GLP-1R and GCGR employ conserved residues to recognize the common segments of their endogenous peptides and use non-conserved residues to make specific interaction that govern the ligand selectivity.

## Discussion

As one of the incretin hormones, GIP modulates glucose metabolism by stimulating the β-cells to release insulin (*29*). Unlike GLP-1, it does not suppress gastric emptying and appetite, while exerting opposite actions on pancreatic α-cells as well as adipocytes leading to glucagon secretion and lipogenesis (*29*). Coupled with reduced sensitivity in type 2 diabetic patients, development of GIPR-based therapeutics met little success (*30*).

Comparison of the full-length structures of six glucagon subfamily of GPCRs demonstrates that bound peptides (GLP-1, exendin-P5, glucagon, ZP3780, secretin, GHRH, GLP-2, and GIP) all adopt a single straight helix with their N-terminus inserted into the TMD core, while the C-terminal is recognized by the ECD (*11, 12, 14, 15, 28*). For parathyroid hormone subfamily of GPCRs, the long-acting PTH analog (LA-PTH) predominantly exhibits an extended helix with its N-terminus inserted deeply into the TMD, where the peptide C-terminus may bend occasionally (*16*). In the case of corticotropin-releasing factor (CRF) subfamily of GPCRs, the N-terminus (first seven residues) of urocortin 1 (UCN1) and CRF1 present an extended loop conformation, and its C-terminal residues (8–40) adopt a single extended helix *(17, 31)*. As far as calcitonin subfamily of GPCRs is concerned, calcitonin gene-related peptide (CGRP) has an unstructured loops in both N- and C-terminal regions (*32*). Looking at pituitary adenylate-cyclase-activating peptide (PACAP) and vasoactive intestinal polypeptide (VIP) receptor subfamily, PACAP displays an extended α-helix, while maxadilan, a natural PAC1R agonist (61-amino acid long), forms the N- and C-terminal helices that are linked as a loop (*10, 31, 33*). These observations highlight diversified peptide binding modes among class B1 GPCRs (Fig. S7).

Species differences in class B1 receptor responsiveness are diversified and receptor-specific, which is tolerable for some receptors such as GLP-1R and GCGR, but leads to concerns for others like GIPR and PTH2R (*34, 35*). Interestingly, the sequence identities between human and mouse at both ligand and receptor levels are more conserved between GLP and GLP-1R (100% and 93%) than that between GIP and GIPR (92% and 81%)(*34*). Such a divergence is not caused by changes in peptide potency, but resides in the biological property of either GIP or the receptor (*34, 36*). However, it may affect GIP-related pharmacology markedly (*34*). Indeed, a previous study found that human GIP is a comparatively weak partial agonist in rodent models (*34*). Human (Pro3)GIP is a full agonist with identical maximum response as human GIP, whereas both rat and mouse (Pro3)GIPs are partial agonists (*34, 36*). Of note is that among rat, mouse and human GIPs, the only residue change (from His to Arg) occurs at the 18^th^ position (*34*). From a structural biology perspective, the variation in the sequences of both GIP (H^18P^/R/R for human, rat and mouse) and GIPR, and the consequent alterations in either peptide-binding or G protein-coupling may offer an explanation. Nonetheless, it may also complicate knowledge transfer from rodents to humans for clinical development of GIPR-based therapeutics.

The interactions between three receptors (GIPR, GLP-1R and GCGR) and their endogenous peptides transduce precise cellular signals responsible for glucose control. This is instituted by a common and closely related mechanism where the extracellular portion of the receptor mainly binds to a cognate ligand while the TMD activates a cascade of signaling events. The upper half of the TMD pocket composed of the top parts of ECL1, TM1 and TM2 interacts with unique residues in the peptide through flexible movement of ECL1 and complementary shape formation by TM1 and TM2, thereby conferring selectively and discriminating unrelated ligands. The lower half of the TMD pocket composed of TMs 3, 6 and 7 displays conserved sequences for recognition of common residues in the peptide. Its key function is to converge external signal into the cytoplasm and executes transduction with high efficiency. This mechanistic design reflects evolutionary advantages because multiple polypeptides could be accurately recognized via different sequences in the upper half of the TMD pocket.

Finally, GIPR, combined with GLP-1R and GCGR, have been intensively studied as targets of dual- or tri- agonists (*37, 38*). Combined activation of GLP-1R and GIPR by dual agonists would provide synergistic and improved effects in glycemic and body weight control (*39*). The GLP-1R/GIPR dual-agonists LY3298176 (developed by Eli Lilly) and NN9709 (developed by Novo Nordisk/Marcadia) as well as GLP-1R/GCGR/GIPR tri-agonist HM15211 (developed by Hamni Pharmaceuticals) are undergoing phase II or III clinical trials (*6*). The detailed structural information on GIPR reported here will certainly be of value to better understand the mode of actions of these therapeutic peptides.

## Materials and Methods

### Cell culture

*Spodoptera frugiperda* (*Sf*9) (Invitrogen) and High-Five™ insect cells (ThermoFisher Scientific) were cultured in ESF 921 serum-free medium (Expression Systems) at 27 C and 120 rpm.

### Constructs

The human GIPR DNA (Genewiz) with one mutation (T345F) was cloned into a modified pFastBac vector (Invitrogen). The native signal peptide was replaced by the haemagglutinin signal peptide (HA) to enhance receptor expression. A BRIL fusion protein was added at the N terminal of the ECD with a TEV protease site and 2GSA linker between them. Forty-five amino acids (Q422-C466) were truncated at the C terminus where LgBiT was added with a 15-amino acid (15AA) polypeptide linker in between, followed by a TEV protease cleavage site and an optimized maltose binding protein- maltose binding protein tag (OMBP-MBP). A dominant-negative bovine Gαs (DNGα_s_)(S54N, G226A, E268A, N271K, K274D, R280K, T284D and I285T) construct was used to stabilize the complex (*11, 40*). SmBiT34 (peptide 86, Promega) subunit was added to the C terminus of Rat Gβ1 with a 15AA polypeptide linker between them. The modified rat Gβ1 and bovine Gγ2 were both cloned into a pFastBac vector.

### Protein expression

Baculoviruses containing the above complex construct were prepared by the Bac-to-Bac system (Invitrogen). GIPR and G_s_ heterotrimer were co-expressed in High-Five™ cells. Briefly, insect cells were grown in ESF 921 culture medium (Expression Systems) to a density of 3.2 × 10^6^ cells/mL, and then cells were infected with four kinds of viral preparations: BRIL-TEV-2GSA-GIPR(22-421)T345F-15AA-LgBiT-TEV-OMBP-MBP, Gα_s_, Gβ1-peptide 86, and Gγ2 at a ratio of 1:3:3:3. After 48 h incubation at 27 C, the cells were collected by centrifugation and stored at −80 C until use.

### Nb35 expression and purification

Nanobody-35 (Nb35) with a 6× his tag at the C terminal was expressed in the periplasm of *E. coli* BL21 (DE3) cells. Briefly, Nb35 target gene was transformed in the bacterium and amplified in TB culture medium with 100 μg/mL ampicillin, 2 mM MgCl_2_, 0.1 % (w/v) glucose at 37 C, 180 rpm. When OD600 reached 0.7-1.2, 1 mM IPTG was added to induce expression followed by overnight incubation at 28 C. The cell pellet was then collected at 3000 rpm under 4 C and stored at −80 C. Nb35 was purified as by size-exclusion chromatography using a HiLoad 16/600 Superdex 75 column (GE Healthcare) with running buffer containing 20 mM HEPES, 100 mM NaCl, pH 7.4. Fractions of Nb35 were concentrated to ~3 mg/mL and quickly frozen in the liquid nitrogen with 10% glycerol and stored in −80 C.

### Complex formation and purification

Cell pellets were lysed in a buffer consisting of 20 mM HEPES, 100 mM NaCl, pH 7.4, 10 mM MgCl_2_, 1 mM MnCl_2_ and 10% glycerol supplemented with protease inhibitor cocktail, EDTA-free (TragetMol). Subsequently, cell membranes were collected by ultracentrifugation at 4 C, 90,000 *g* for 35 min. The membranes were resuspended with a buffer containing 20 mM HEPES, 100 mM NaCl, pH 7.4, 10 mM MgCl_2_, 1 mM MnCl_2_ and 10% glycerol. The complex of GIPR-G_s_ was assembled by adding 15 μM GIP_1-42_ (GenScript), 100 μM TCEP, 25 mU/mL Apyrase (Sigma-Aldrich), 15 μg/mL Nb35 and 100 U salt active nuclease (Sigma-Aldrich) supplemented with protease inhibitor cocktail for 1.5 h incubation at room temperature (RT). The preparation was then solubilized with 0.5% (w/v) lauryl maltose neopentylglycol (LMNG, Anatrace) and 0.1% (w/v) cholesterol hemisuccinate (CHS, Anatrace) with additional 1 μM GIP_1-42_ for 3 h at 4 C. The supernatant was isolated by centrifugation at 90,000 *g* for 35 min and the solubilized complex was incubated with amylose resin (NEB) for 2.5 h at 4 C. After batch binding, the resin was collected by centrifugation at 550 *g* and loaded onto a gravity flow column. The resin in column was firstly washed with 5 column volumes of buffer containing 20 mM HEPES, pH 7.4, 100 mM NaCl, 10% (v/v) glycerol, 5 mM MgCl_2_, 1 mM MnCl_2_, 25 μM TCEP, 3 μM GIP_1-42_, 0.1% (w/v) LMNG and 0.02% (w/v) CHS. Subsequently, the resin was washed with 25 column volumes of buffer containing 20 mM HEPES, pH 7.4, 100 mM NaCl, 10% (v/v) glycerol, 5 mM MgCl_2_, 1 mM MnCl_2_, 25 μM TCEP, 3 μM GIP_1-42_, 0.03% (w/v) LMNG, 0.01% (w/v) glyco-diosgenin (GDN, Anatrace) and 0.008% (w/v) CHS. The protein was then incubated with a buffer containing 20 mM HEPES, pH 7.4, 100 mM NaCl, 10% (v/v) glycerol, 5 mM MgCl_2_, 1 mM MnCl_2_, 25 μM TCEP, 50 μM GIP_1-42_, 10 μg/mL Nb35, 0.03% (w/v) LMNG, 0.01% (w/v) glyco-diosgenin, 0.008% (w/v) CHS and 30 μg/mL His-tagged TEV protease on the column overnight at 4 C. The flow through was collected and concentrated to 500 μL using a 100 kDa filter (Merck Millipore). Size-exclusion chromatography was performed by loading the protein onto Superose 6 Increase 10/300GL (GE Healthcare) column with running buffer containing 20 mM HEPES, pH 7.4, 100 mM NaCl, 10 mM MgCl_2_, 100 μM TCEP, 5 μM GIP_1-42_, 0.00075% (w/v) LMNG, 0.00025% (w/v) glyco-diosgenin, 0.0002% (w/v) CHS and 0.00025% digitonin (Anatrace). Monomeric GIPR-G_s_ complexes were collected and concentrated for cryo-EM analysis.

### Data acquisition and image processing

The purified GIP_1-42_–GIPR–G_s_–Nb35 complex at a concentration of 6-7 mg/mL was mixed with 100 μM GIP_1-42_ at 4 C and applied to glow-discharged holey carbon grids (Quantifoil R1.2/1.3, Au 300 mesh) that were subsequently vitrified by plunging into liquid ethane using a Vitrobot Mark IV (ThermoFisher Scientific). A Titan Krios equipped with a Gatan K3 Summit direct electron detector was used to acquire Cryo-EM images. The microscope was operated at 300 kV accelerating voltage, at a nominal magnification of 46685× in counting mode, corresponding to a pixel size of 1.071Å. Totally, 8023 movies were obtained with a defocus range of −1.2 to −2.2 μm. An accumulated dose of 80 electrons per Å^2^ was fractionated into a movie stack of 36 frames.

Dose-fractionated image stacks were subjected to beam-induced motion correction using MotionCor2.1. A sum of all frames, filtered according to the exposure dose, in each image stack was used for further processing. Contrast transfer function parameters for each micrograph were determined by Gctf v1.06. Particle selection, 2D and 3D classifications were performed on a binned dataset with a pixel size of 2.142 Å using RELION-3.0-beta2. Auto-picking yielded 4,895,399 particle projections that were subjected to reference-free 2D classification to discard false positive particles or particles categorized in poorly defined classes, producing 2,754,623 particle projections for further processing. This subset of particle projections was subjected to a round of maximum-likelihood-based three dimensional classifications with a pixel size of 2.142 Å, resulting in one well-defined subset with 1,395,031 projections. Further 3D classifications with mask on the receptor produced one good subset accounting for 565,239 particles, which were subjected to another round of 3D classifications with mask on the ECD. A selected subset containing 295,021 projections was then subjected to 3D refinement and Bayesian polishing with a pixel size of 1.071 Å. After the last round of refinement, the final map has an indicated global resolution of 2.94 Å at a Fourier shell correlation (FSC) of 0.143. Local resolution was determined using the Bsoft package with half maps as input maps.

### Model building and refinement

The cryo-EM structure of GCGR–G_s_–Nb35 complex (PDB code 6WPW)(*14*) and the crystal structure (PDB code 2QKH) (*21*) were used as the start for model building and refinement against the EM map. The model was docked into the EM density map using Chimera (*41*), followed by iterative manual adjustment and rebuilding in COOT (*42*). Real space refinement was performed using Phenix (*43*). The model statistics were validated using MolProbity (*44*). Structural figures were prepared in Chimera and PyMOL (https://pymol.org/2/). The final refinement statistics are provided in Table S1.

### cAMP accumulation assay

GIP_1-42_ stimulated cAMP accumulation was measured by a LANCE Ultra cAMP kit (PerkinElmer). Briefly, HEK 293T cells were cultured in DMEM (Gibco) supplemented with 10% (v/v) fetal bovine serum (FBS, Gibco) and 1% (v/v) sodium pyruvate (Gibco) at 37 C, 5% CO_2_. Cells were seeded onto 6-well cell culture plates and transiently transfected with different GIPR constructs using Lipofectamine 2000 transfection reagent (Invitrogen). All the mutant constructs were modified by single-point mutation in the setting of the WT construct (HA-Flag-3GSA-GIPR(22-466)). After 24 h culture, the transfected cells were seeded onto 384-well microtiter plates at a density of 3000 cells per well in HBSS supplemented with 5 mM HEPES, 0.1% (w/v) bovine serum albumin (BSA) and 0.5 mM 3-isobutyl-1-methylxanthine. The cells were stimulated with different concentrations of GIP_1-42_ for 40 min at RT. Eu and Ulight were then diluted by cAMP detection buffer and added to the plates separately to terminate the reaction. Plates were incubated at RT for 40 min and the fluorescence intensity measured at 620 nm and 650 nm by an EnVision multilabel plate reader (PerkinElmer).

### Whole cell binding assay

CHO-K1 cells were cultured in F12 medium with 10% FBS and seeded at a density of 30,000 cells/well in Isoplate-96 plates (PerkinElmer). The WT (HA-Flag-3GSA-GIPR(22-466)) or mutant GIPR were transiently transfected using Lipofectamine 2000 transfection reagent. The mutant construct was modified by single-point mutation in the setting of the WT construct. Twenty-four hours after transfection, cells were washed twice, and incubated with blocking buffer (F12 supplemented with 33 mM HEPES and 0.1% bovine serum albumin (BSA), pH 7.4) for 2 h at 37°C. For homogeneous binding, cells were incubated in binding buffer with a constant concentration of ^125^I-GIP (40 pM, PerkinElmer) and increasing concentrations of unlabeled GIP_1-42_ (3.57 pM to 1 μM) at RT for 3 h. Following incubation, cells were washed three times with ice-cold PBS and lysed by addition of 50 μL lysis buffer (PBS supplemented with 20 mM Tris-HCl, 1% Triton X-100, pH 7.4). Fifty µL of scintillation cocktail (OptiPhase SuperMix, PerkinElmer) was added and the plates were subsequently counted for radioactivity (counts per minute, CPM) in a scintillation counter (MicroBeta2 Plate Counter, PerkinElmer).

### Receptor surface expression

Cell surface expression was determined by flow cytometry to the N-terminal Flag tag on the WT GIPR (HA-Flag-3GSA-GIPR(22-466)) and its mutants transiently expressed in HEK 293T cells. All the mutant constructs were modified by single-point mutation in the setting of the WT construct. Briefly, approximately 2 × 10^5^ cells were blocked with PBS containing 5% BSA (w/v) at RT for 15 min, and then incubated with 1:300 anti-Flag primary antibody (diluted with PBS containing 5% BSA, Sigma-Aldrich) at RT for 1 h. The cells were then washed three times with PBS containing 1% BSA (w/v) followed by 1 h incubation with 1:1000 anti-mouse Alexa Fluor 488 conjugated secondary antibody (diluted with PBS containing 5% BSA, Invitrogen) at RT in the dark. After washing three times, cells were re-suspended in 200 μL PBS containing 1% BSA for detection by NovoCyte (Agilent) utilizing laser excitation and emission wavelengths of 488 nm and 530 nm, respectively. For each sample, 20,000 cellular events were collected, and the total fluorescence intensity of positive expression cell population was calculated. Data were normalized to the WT receptor.

### Molecular dynamics simulations

Molecular dynamic simulations were performed by Gromacs 2018.5. The peptide–GIPR complexes were built based on the cryo-EM GIP–GIPR–G_s_ complex and prepared by the Protein Preparation Wizard (Schrodinger 2017-4) with the G protein and Nb35 nanobody removed. The receptor chain termini were capped with acetyl and methylamide, and the titratable residues were left in their dominant state at pH 7.0. The complexes were embedded in a bilayer composed of 200 POPC lipids and solvated with 0.15 M NaCl in explicitly TIP3P waters using CHARMM-GUI Membrane Builder (*45*). The CHARMM36-CAMP force filed (*46*) was adopted for protein, peptides, lipids and salt ions. The Particle Mesh Ewald (PME) method was used to treat all electrostatic interactions beyond a cut-off of 10Å and the bonds involving hydrogen atoms were constrained using LINCS algorithm (*47*). The complex system was firstly relaxed using the steepest descent energy minimization, followed by slow heating of the system to 310 K with restraints. The restraints were reduced gradually over 50 ns. Finally, restrain-free production run was carried out for each simulation, with a time step of 2 fs in the NPT ensemble at 310 K and 1 bar using the Nose-Hoover thermostat and the semi-isotropic Parrinello-Rahman barostat (*48*), respectively. The buried interface areas were calculated with FreeSASA (*49*) using the Sharke-Rupley algorithm with a probe radius of 1.2 Å. The last 700 ns trajectory of each simulation was used to root mean square fluctuation (RMSF) calculation.

### Statistical analysis

All functional data were presented as means ± standard error of the mean (SEM). Statistical analysis was performed using GraphPad Prism 7 (GraphPad Software). Concentration-response curves were evaluated with a three-parameter logistic equation. The significance was determined with either two-tailed Student’s *t*-test or one-way ANOVA. Significant difference is accepted at P < 0.05.

## Acknowledgments

We thank Elita Yuliantie, Wen Sun, Zhaotong Cong, Fulai Zhou, Yuqi Ping, X. Edward Zhou, Karsten Melcher, Jinhuan Chen and Xijiang Pan for technical advice. The cryo-EM data were collected at Cryo-Electron Microscopy Research Center, Shanghai Institute of Materia Medica. This work was partially supported by National Natural Science Foundation of China 81872915 (M.-W.W.), 32071203 (L.H.Z), 81773792 (D.H.Y.), 81973373 (D.H.Y.) and 21704064 (Q.T.Z.); National Science and Technology Major Project of China – Key New Drug Creation and Manufacturing Program 2018ZX09735–001 (M.-W.W.) and 2018ZX09711002–002–005 (D.H.Y.); National Key Basic Research Program of China 2018YFA0507000 (M.-W.W.); Ministry of Science and Technology of China 2018YFA0507002 (H.E.X.); Shanghai Municipal Science and Technology Major Project 2019SHZDZX02 (H.E.X.); Strategic Priority Research Program of Chinese Academy of Sciences XDB37030103 (H.E.X.); Shanghai Municipality Science and Technology Development Fund 18430711500 (M.-W.W.);Novo Nordisk-CAS Research Fund grant NNCAS-2017–1-CC (D.H.Y.); Shanghai Science and Technology Development Fund 18ZR1447800 (L.H.Z.); The Young Innovator Association of CAS 2018325 (L.H.Z.) and SA-SIBS Scholarship Program (L.H.Z. and D.H.Y.)

## Author contributions

F.H.Z., C.Z., L.H.Z. and K.N.H. designed the expression constructs, purified the receptor complexes, prepared the final samples for negative stain/data collection towards the structure and participated in manuscript preparation; X.Y.Z. performed cryo-EM map calculation and figure preparation. Q.T.Z. and L.H.Z. performed structural analysis; Q.T.Z. conducted MD simulations, figure preparation, and participated in manuscript writing; A.T.D. and Y.C. conducted ligand binding and signaling experiments; W.C.Y. and Q.D.R. assisted in complex purification; D.D.S. and Y.Z. helped cryo-EM sample preparation; F.W. and T.X. took part in structural analysis; D.H.Y. supervised mutagenesis and signaling experiment; R.C.S., H.E.X. and M.-W.W. initiated the project; H.E.X., D.H.Y. L.H.Z. and M.-W.W. supervised the studies, analyzed the data and wrote the manuscript with inputs from all co-authors.

## Data and materials availability

All data are available in the main text or the supplementary materials.

## References

1. S. J. Kim, C. Nian, C. H. McIntosh, Activation of lipoprotein lipase by glucose-dependent insulinotropic polypeptide in adipocytes. A role for a protein kinase B, LKB1, and AMP-activated protein kinase cascade. J Biol Chem 282, 8557–8567 (2007).

2. E. Faivre, C. Holscher, Neuroprotective effects of D-Ala(2)GIP on Alzheimer’s disease biomarkers in an APP/PS1 mouse model. Alzheimers Res Ther 5, 20 (2013).

3. M. C. Lagerstrom, H. B. Schioth, Structural diversity of G protein-coupled receptors and significance for drug discovery. Nat Rev Drug Discov 7, 339–357 (2008).

4. B. Finan, C. Clemmensen, Z. Zhu, K. Stemmer, K. Gauthier, L. Muller, M. De Angelis, K. Moreth, F. Neff, D. Perez-Tilve, K. Fischer, D. Lutter, M. A. Sanchez-Garrido, P. Liu, J. Tuckermann, M. Malehmir, M. E. Healy, A. Weber, M. Heikenwalder, M. Jastroch, M. Kleinert, S. Jall, S. Brandt, F. Flamant, K. W. Schramm, H. Biebermann, Y. Doring, C. Weber, K. M. Habegger, M. Keuper, V. Gelfanov, F. Liu, J. Kohrle, J. Rozman, H. Fuchs, V. Gailus-Durner, M. Hrabe de Angelis, S. M. Hofmann, B. Yang, M. H. Tschop, R. DiMarchi, T. D. Muller, Chemical hybridization of glucagon and thyroid hormone optimizes therapeutic impact for metabolic disease. Cell 167, 843–857 e814 (2016).

5. C. Longuet, E. M. Sinclair, A. Maida, L. L. Baggio, M. Maziarz, M. J. Charron, D. J. Drucker, The glucagon receptor is required for the adaptive metabolic response to fasting. Cell Metab 8, 359–371 (2008).

6. D. Yang, Q. Zhou, V. Labroska, S. Qin, S. Darbalaei, Y. Wu, E. Yuliantie, L. Xie, H. Tao, J. Cheng, Q. Liu, S. Zhao, W. Shui, Y. Jiang, M. W. Wang, G protein-coupled receptors: structure- and function-based drug discovery. Signal Transduct Target Ther 6, 7 (2021).

7. C. Parthier, S. Reedtz-Runge, R. Rudolph, M. T. Stubbs, Passing the baton in class B GPCRs: peptide hormone activation via helix induction? Trends Biochem Sci 34, 303–310 (2009).

8. C. M. Koth, J. M. Murray, S. Mukund, A. Madjidi, A. Minn, H. J. Clarke, T. Wong, V. Chiang, E. Luis, A. Estevez, J. Rondon, Y. Zhang, I. Hotzel, B. B. Allan, Molecular basis for negative regulation of the glucagon receptor. Proc Natl Acad Sci U S A 109, 14393–14398 (2012).

9. L. Yang, D. Yang, C. de Graaf, A. Moeller, G. M. West, V. Dharmarajan, C. Wang, F. Y. Siu, G. Song, S. Reedtz-Runge, B. D. Pascal, B. Wu, C. S. Potter, H. Zhou, P. R. Griffin, B. Carragher, H. Yang, M. W. Wang, R. C. Stevens, H. Jiang, Conformational states of the full-length glucagon receptor. Nat Commun 6, 7859 (2015).

10. J. Duan, D. D. Shen, X. E. Zhou, P. Bi, Q. F. Liu, Y. X. Tan, Y. W. Zhuang, H. B. Zhang, P. Y. Xu, S. J. Huang, S. S. Ma, X. H. He, K. Melcher, Y. Zhang, H. E. Xu, Y. Jiang, Cryo-EM structure of an activated VIP1 receptor-G protein complex revealed by a NanoBiT tethering strategy. Nat Commun 11, 4121 (2020).

11. F. Zhou, H. Zhang, Z. Cong, L. H. Zhao, Q. Zhou, C. Mao, X. Cheng, D. D. Shen, X. Cai, C. Ma, Y. Wang, A. Dai, Y. Zhou, W. Sun, F. Zhao, S. Zhao, H. Jiang, Y. Jiang, D. Yang, H. Eric Xu, Y. Zhang, M. W. Wang, Structural basis for activation of the growth hormone-releasing hormone receptor. Nat Commun 11, 5205 (2020).

12. W. Sun, L. N. Chen, Q. Zhou, L. H. Zhao, D. Yang, H. Zhang, Z. Cong, D. D. Shen, F. Zhao, F. Zhou, X. Cai, Y. Chen, Y. Zhou, S. Gadgaard, W. J. C. van der Velden, S. Zhao, Y. Jiang, M. M. Rosenkilde, H. E. Xu, Y. Zhang, M. W. Wang, A unique hormonal recognition feature of the human glucagon-like peptide-2 receptor. Cell Res 30, 1098–1108 (2020).

13. D. Wootten, J. Simms, L. J. Miller, A. Christopoulos, P. M. Sexton, Polar transmembrane interactions drive formation of ligand-specific and signal pathway-biased family B G protein-coupled receptor conformations. Proc Natl Acad Sci U S A 110, 5211–5216 (2013).

14. A. Qiao, S. Han, X. Li, Z. Li, P. Zhao, A. Dai, R. Chang, L. Tai, Q. Tan, X. Chu, L. Ma, T. S. Thorsen, S. Reedtz-Runge, D. Yang, M. W. Wang, P. M. Sexton, D. Wootten, F. Sun, Q. Zhao, B. Wu, Structural basis of Gs and Gi recognition by the human glucagon receptor. Science 367, 1346–1352 (2020).

15. Y. Zhang, B. Sun, D. Feng, H. Hu, M. Chu, Q. Qu, J. T. Tarrasch, S. Li, T. Sun Kobilka, B. K. Kobilka, G. Skiniotis, Cryo-EM structure of the activated GLP-1 receptor in complex with a G protein. Nature 546, 248–253 (2017).

16. L. H. Zhao, S. Ma, I. Sutkeviciute, D. D. Shen, X. E. Zhou, P. W. de Waal, C. Y. Li, Y. Kang, L. J. Clark, F. G. Jean-Alphonse, A. D. White, D. Yang, A. Dai, X. Cai, J. Chen, C. Li, Y. Jiang, T. Watanabe, T. J. Gardella, K. Melcher, M. W. Wang, J. P. Vilardaga, H. E. Xu, Y. Zhang, Structure and dynamics of the active human parathyroid hormone receptor-1. Science 364, 148–153 (2019).

17. S. Ma, Q. Shen, L. H. Zhao, C. Mao, X. E. Zhou, D. D. Shen, P. W. de Waal, P. Bi, C. Li, Y. Jiang, M. W. Wang, P. M. Sexton, D. Wootten, K. Melcher, Y. Zhang, H. E. Xu, Molecular basis for hormone recognition and activation of corticotropin-releasing factor receptors. Mol Cell 77, 669–680 e664 (2020).

18. T. Yaqub, I. G. Tikhonova, J. Lattig, R. Magnan, M. Laval, C. Escrieut, C. Boulegue, C. Hewage, D. Fourmy, Identification of determinants of glucose-dependent insulinotropic polypeptide receptor that interact with N-terminal biologically active region of the natural ligand. Mol Pharmacol 77, 547–558 (2010).

19. B. D. Kerr, A. J. Flatt, P. R. Flatt, V. A. Gault, Characterization and biological actions of N-terminal truncated forms of glucose-dependent insulinotropic polypeptide. Biochem. Biophys. Res. Commun 404, 870–876 (2011).

20. M. B. N. Gabe, W. J. C. van der Velden, F. X. Smit, L. S. Gasbjerg, M. M. Rosenkilde, Molecular interactions of full-length and truncated GIP peptides with the GIP receptor - A comprehensive review. Peptides 125, 170224 (2020).

21. C. Parthier, M. Kleinschmidt, P. Neumann, R. Rudolph, S. Manhart, D. Schlenzig, J. Fanghanel, J. U. Rahfeld, H. U. Demuth, M. T. Stubbs, Crystal structure of the incretin-bound extracellular domain of a G protein-coupled receptor. Proc Natl Acad Sci U S A 104, 13942–13947 (2007).

22. D. Hilger, K. K. Kumar, H. Hu, M. F. Pedersen, E. S. O’Brien, L. Giehm, C. Jennings, G. Eskici, A. Inoue, M. Lerch, J. M. Mathiesen, G. Skiniotis, B. K. Kobilka, Structural insights into differences in G protein activation by family A and family B GPCRs. Science 369, (2020).

23. A. Jazayeri, A. S. Dore, D. Lamb, H. Krishnamurthy, S. M. Southall, A. H. Baig, A. Bortolato, M. Koglin, N. J. Robertson, J. C. Errey, S. P. Andrews, I. Teobald, A. J. Brown, R. M. Cooke, M. Weir, F. H. Marshall, Extra-helical binding site of a glucagon receptor antagonist. Nature 533, 274–277 (2016).

24. H. Zhang, A. Qiao, L. Yang, N. Van Eps, K. S. Frederiksen, D. Yang, A. Dai, X. Cai, H. Zhang, C. Yi, C. Cao, L. He, H. Yang, J. Lau, O. P. Ernst, M. A. Hanson, R. C. Stevens, M. W. Wang, S. Reedtz-Runge, H. Jiang, Q. Zhao, B. Wu, Structure of the glucagon receptor in complex with a glucagon analogue. Nature 553, 106–110 (2018).

25. R. Chang, X. Zhang, A. Qiao, A. Dai, M. J. Belousoff, Q. Tan, L. Shao, L. Zhong, G. Lin, Y. L. Liang, L. Ma, S. Han, D. Yang, R. Danev, M. W. Wang, D. Wootten, B. Wu, P. M. Sexton, Cryo-electron microscopy structure of the glucagon receptor with a dual-agonist peptide. J Biol Chem 295, 9313–9325 (2020).

26. H. Zhang, A. Qiao, D. Yang, L. Yang, A. Dai, C. de Graaf, S. Reedtz-Runge, V. Dharmarajan, H. Zhang, G. W. Han, T. D. Grant, R. G. Sierra, U. Weierstall, G. Nelson, W. Liu, Y. Wu, L. Ma, X. Cai, G. Lin, X. Wu, Z. Geng, Y. Dong, G. Song, P. R. Griffin, J. Lau, V. Cherezov, H. Yang, M. A. Hanson, R. C. Stevens, Q. Zhao, H. Jiang, M. W. Wang, B. Wu, Structure of the full-length glucagon class B G-protein-coupled receptor. Nature 546, 259–264 (2017).

27. X. Zhang, M. J. Belousoff, P. Zhao, A. J. Kooistra, T. T. Truong, S. Y. Ang, C. R. Underwood, T. Egebjerg, P. Senel, G. D. Stewart, Y. L. Liang, A. Glukhova, H. Venugopal, A. Christopoulos, S. G. B. Furness, L. J. Miller, S. Reedtz-Runge, C. J. Langmead, D. E. Gloriam, R. Danev, P. M. Sexton, D. Wootten, Differential GLP-1R binding and activation by peptide and non-peptide agonists. Mol Cell 80, 485–500 e487 (2020).

28. M. Dong, G. Deganutti, S. J. Piper, Y. L. Liang, M. Khoshouei, M. J. Belousoff, K. G. Harikumar, C. A. Reynolds, A. Glukhova, S. G. B. Furness, A. Christopoulos, R. Danev, D. Wootten, P. M. Sexton, L. J. Miller, Structure and dynamics of the active Gs-coupled human secretin receptor. Nat Commun 11, 4137 (2020).

29. Y. Seino, M. Fukushima, D. Yabe, GIP and GLP-1, the two incretin hormones: Similarities and differences. J Diabetes Investig 1, 8–23 (2010).

30. T. Coskun, K. W. Sloop, C. Loghin, J. Alsina-Fernandez, S. Urva, K. B. Bokvist, X. Cui, D. A. Briere, O. Cabrera, W. C. Roell, U. Kuchibhotla, J. S. Moyers, C. T. Benson, R. E. Gimeno, D. A. D’Alessio, A. Haupt, LY3298176, a novel dual GIP and GLP-1 receptor agonist for the treatment of type 2 diabetes mellitus: From discovery to clinical proof of concept. Mol Metab 18, 3–14 (2018).

31. Y. L. Liang, M. J. Belousoff, P. Zhao, C. Koole, M. M. Fletcher, T. T. Truong, V. Julita, G. Christopoulos, H. E. Xu, Y. Zhang, M. Khoshouei, A. Christopoulos, R. Danev, P. M. Sexton, D. Wootten, Toward a structural understanding of class B GPCR peptide binding and activation. Mol Cell 77, 656–668 e655 (2020).

32. Y. L. Liang, M. Khoshouei, G. Deganutti, A. Glukhova, C. Koole, T. S. Peat, M. Radjainia, J. M. Plitzko, W. Baumeister, L. J. Miller, D. L. Hay, A. Christopoulos, C. A. Reynolds, D. Wootten, P. M. Sexton, Cryo-EM structure of the active, Gs-protein complexed, human CGRP receptor. Nature 561, 492–497 (2018).

33. J. Wang, X. Song, D. Zhang, X. Chen, X. Li, Y. Sun, C. Li, Y. Song, Y. Ding, R. Ren, E. H. Harrington, L. A. Hu, W. Zhong, C. Xu, X. Huang, H. W. Wang, Y. Ma, Cryo-EM structures of PAC1 receptor reveal ligand binding mechanism. Cell Res 30, 436–445 (2020).

34. A. H. Sparre-Ulrich, L. S. Hansen, B. Svendsen, M. Christensen, F. K. Knop, B. Hartmann, J. J. Holst, M. M. Rosenkilde, Species-specific action of (Pro3)GIP - a full agonist at human GIP receptors, but a partial agonist and competitive antagonist at rat and mouse GIP receptors. Br J Pharmacol 173, 27–38 (2016).

35. S. R. Hoare, T. I. Bonner, T. B. Usdin, Comparison of rat and human parathyroid hormone 2 (PTH2) receptor activation: PTH is a low potency partial agonist at the rat PTH2 receptor. Endocrinology 140, 4419–4425 (1999).

36. C. J. Bailey, GIP analogues and the treatment of obesity-diabetes. Peptides 125, 170202 (2020).

37. M. A. Skow, N. C. Bergmann, F. K. Knop, Diabetes and obesity treatment based on dual incretin receptor activation: ‘twincretins’. Diabetes Obes Metab 18, 847–854 (2016).

38. K. Alexiadou, O. Anyiam, T. Tan, Cracking the combination: Gut hormones for the treatment of obesity and diabetes. J Neuroendocrinol 31, e12664 (2019).

39. M. Bastin, F. Andreelli, Dual GIP-GLP1-receptor agonists in the treatment of type 2 diabetes: a short review on emerging data and therapeutic potential. Diabetes Metab Syndr Obes 12, 1973–1985 (2019).

40. Y. L. Liang, P. Zhao, C. Draper-Joyce, J. A. Baltos, A. Glukhova, T. T. Truong, L. T. May, A. Christopoulos, D. Wootten, P. M. Sexton, S. G. B. Furness, Dominant negative G proteins enhance formation and purification of agonist-GPCR-G protein complexes for structure determination. ACS Pharmacol Transl Sci 1, 12–20 (2018).

41. E. F. Pettersen, T. D. Goddard, C. C. Huang, G. S. Couch, D. M. Greenblatt, E. C. Meng, T. E. Ferrin, UCSF Chimera--a visualization system for exploratory research and analysis. Journal of computational chemistry 25, 1605–1612 (2004).

42. P. Emsley, K. Cowtan, Coot: model-building tools for molecular graphics. Acta Crystallogr D Biol Crystallogr 60, 2126–2132 (2004).

43. P. D. Adams, P. V. Afonine, G. Bunkoczi, V. B. Chen, I. W. Davis, N. Echols, J. J. Headd, L. W. Hung, G. J. Kapral, R. W. Grosse-Kunstleve, A. J. McCoy, N. W. Moriarty, R. Oeffner, R. J. Read, D. C. Richardson, J. S. Richardson, T. C. Terwilliger, P. H. Zwart, PHENIX: a comprehensive Python-based system for macromolecular structure solution. Acta Crystallogr D Biol Crystallogr 66, 213–221 (2010).

44. V. B. Chen, W. B. Arendall, 3rd, J. J. Headd, D. A. Keedy, R. M. Immormino, G. J. Kapral, L. W. Murray, J. S. Richardson, D. C. Richardson, MolProbity: all-atom structure validation for macromolecular crystallography. Acta Crystallogr D Biol Crystallogr 66, 12–21 (2010).

45. E. L. Wu, X. Cheng, S. Jo, H. Rui, K. C. Song, E. M. Davila-Contreras, Y. Qi, J. Lee, V. Monje-Galvan, R. M. Venable, J. B. Klauda, W. Im, CHARMM-GUI Membrane Builder toward realistic biological membrane simulations. J. Comput. Chem 35, 1997–2004 (2014).

46. O. Guvench, S. S. Mallajosyula, E. P. Raman, E. Hatcher, K. Vanommeslaeghe, T. J. Foster, F. W. Jamison, 2nd, A. D. Mackerell, Jr., CHARMM additive all-atom force field for carbohydrate derivatives and its utility in polysaccharide and carbohydrate-protein modeling. J Chem Theory Comput 7, 3162–3180 (2011).

47. B. Hess, P-LINCS: a parallel linear constraint solver for Molecular Simulation. J Chem Theory Comput 4, 116–122 (2008).

48. K. M. Aoki, F. Yonezawa, Constant-pressure molecular-dynamics simulations of the crystal-smectic transition in systems of soft parallel spherocylinders. Phys Rev A 46, 6541–6549 (1992).

49. S. Mitternacht, FreeSASA: An open source C library for solvent accessible surface area calculations. F1000Res 5, 189 (2016).

